# Spleen glia are a transcriptionally unique glial subtype interposed between immune cells and sympathetic axons

**DOI:** 10.1101/2020.10.12.336446

**Authors:** Tawaun A. Lucas, Li Zhu, Marion S. Buckwalter

## Abstract

Glia are known to play important roles in the brain, the gut, and around the sciatic nerve. While the gut has its own specialized nervous system, other viscera are innervated solely by autonomic nerves. The functions of glia that accompany autonomic innervation are not well known, even though they are one of the most abundant cell types in the peripheral nervous system. Here, we focused on non-myelinating Schwann Cells in the spleen, spleen glia. The spleen is a major immune organ innervated by the sympathetic nervous system, which modulates immune function. This interaction is known as neuroimmune communication. We establish that spleen glia can be visualized using both immunohistochemistry for S100B and GFAP and with a reporter mouse. Spleen glia ensheath sympathetic axons and are localized to the lymphocyte-rich white pulp areas of the spleen. We sequenced the spleen glia transcriptome and identified genes that are likely involved in axonal ensheathment and communication with both nerves and immune cells. Spleen glia express receptors for neurotransmitters made by sympathetic axons (adrenergic, purinergic, and Neuropeptide Y), and also cytokines, chemokines, and their receptors that may communicate with immune cells in the spleen. We also established similarities and differences between spleen glia and other glial types. While all glia share many genes in common, spleen glia differentially express immune genes, including genes involved in cytokine-cytokine receptor interactions, phagocytosis, and the complement cascade. Thus, spleen glia are a unique glial type, physically and transcriptionally poised to participate in neuroimmune communication in the spleen.

**Table of Contents:** 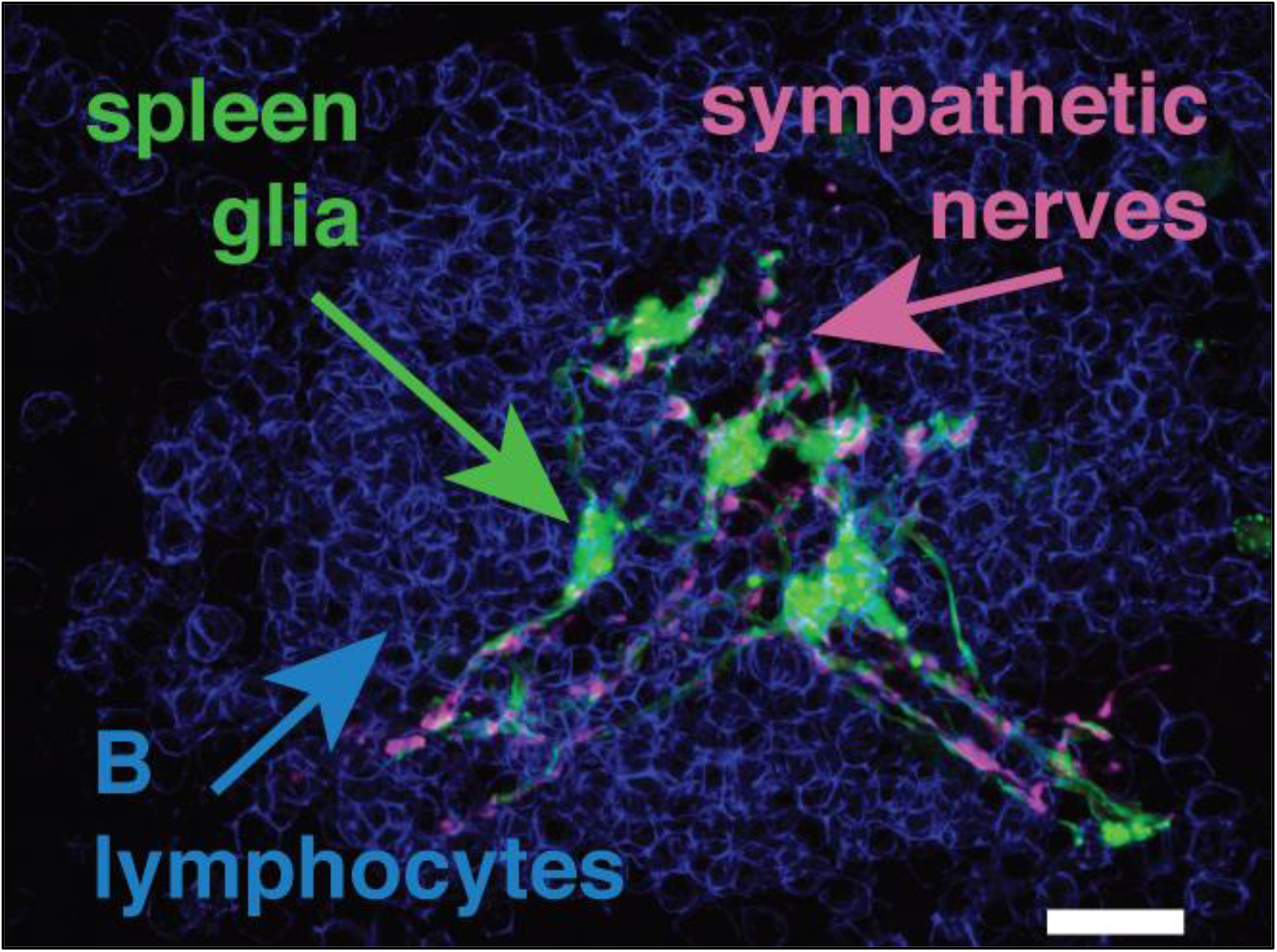

**Main Points:**

- Spleen glia maintain tight associations with splenic nerves and come in close contact with immune cells
- Spleen glia express genes required for communication with nerves and immune cells
- Spleen glia are a transcriptionally unique glial type

## Introduction

Glia play important and complex functional roles in both the central (CNS) and peripheral (PNS) nervous systems. In the brain, astrocytes integrate and process synaptic information and control synaptic transmission and plasticity (Perea et al., 2009). On the other hand, enteric glia in the gut modulate gut motility via bidirectional signaling with neurons, altering smooth muscle cell contraction (Grubišić et al., 2018). While the gut has its own specialized nervous system, other visceral organs are innervated solely by autonomic (sympathetic and parasympathetic) nerves. The functions of non-myelinating glia that accompany autonomic innervation are not well known, despite being one of the most abundant cell types in the PNS (Griffin & Thompson, 2008). Although the structure and function of glia in each visceral organ is relatively unknown, they are likely specialized to the nerves they accompany and the organs in which they reside. This may be particularly important for glia in immune organs such as the spleen, in which the nerves ensheathed by glia come in close contact with immune cells (Felten et al., 1987), potentially positioning them to contribute to neuroimmune crosstalk.

Non-myelinating Schwann cells in the spleen (spleen glia) associate with axons and are derived from the neural crest (Barlow-Anacker et. al. 2017; Felten et al., 1987; Heusermann & Stutte, 1977). The spleen is the largest secondary lymphoid organ and plays a vital role in detecting bloodborne antigens and conferring immunologic memory. It is innervated by the splenic nerve which is predominantly sympathetic. The sympathetic nervous system modulates immunity (Bellinger et al., 2008; Bucsek et al., 2018; Deng et al., 2004; Miller et al., 2019), and epinephrine and norepinephrine released from the adrenal glands and sympathetic nerves within organs act on adrenergic receptors expressed by immune cells (Klehmet et al., 2009; Miller et al., 2019; Pruss et al., 2017). An example of this in the spleen occurs after stroke, when the sympathetic nervous system mediates the death of B-lymphocytes, thus increasing the susceptibility to infection (McCulloch et al., 2017). Given the well-described functional roles of other glia such as altering neurotransmission, releasing cytokines and chemokines, and phagocytosis, spleen glia may serve to modulate this and other neuroimmune interactions in the spleen. However, a complete characterization of their localization within the organ and their anatomical interactions with other cell types is lacking. Additionally, the genomic identity of spleen glia is unknown. A comprehensive anatomical and transcriptomic description of spleen glia is therefore essential to determine if they are localized near immune cells, whether they express receptors and other genes that might be used for neuroimmune crosstalk in the spleen, determine what other processes they may mediate, and to begin to understand how similar they are to other glial types.

Thus, in our research we utilized immunohistochemistry and spleen glia reporter mice to characterize the anatomy and morphology of spleen glia, including their anatomical interactions with sympathetic axons and immune cells. In addition, we generated a transcriptome of spleen glia to define which mRNAs they express and infer possible functions. Lastly, we compared the transcriptome of spleen glia to that of other CNS and PNS glia to determine similarities and differences between different glial cell types.

## Materials and Methods

### Animals

All animal procedures were performed in accordance with the protocol approved by the Institutional Animal Care and Use Committee at Stanford University. C57BL/6J mice aged 8 to 12-weeks were purchased from The Jackson Laboratory, Bar Harbor, ME, and utilized for all studies: GFAP-Cre (JAX#12886) and Rosa26 eGFP (JAX#4077).

### Perfusion and tissue preparation

Mice were anesthetized with 6% chloral hydrate in PBS and terminally perfused through the left ventricle with 10–20ml of cold 0.9% heparinized saline (10 units/ml). Spleens were collected and fixed in 4% paraformaldehyde in phosphate buffer overnight then transferred to 30% sucrose in phosphate buffer until sunken, usually overnight. PFA-fixed spleens were sectioned in both sagittal and coronal planes using a freezing sliding microtome to generate 40 μm thick sections (Microm HM430).

### Immunohistochemistry and antibodies

Immunohistochemistry was performed on PFA-fixed free-floating tissue sections using standard protocols. Sections were washed in Tris Buffered Saline (TBS) and blocked with 5% serum for 1 hour. Primary antibodies were diluted in 0.1% Triton X-100 and 3% serum and applied overnight at room temperature. We used the following primary antibodies: rabbit anti-GFAP at 1:3,000 (Dako), rabbit anti-S100B at 1:500 (Dako), rabbit anti-TH at 1:500 (Millipore), rabbit anti-Pgp9.5 at 1:500 (Cedarlane), hamster anti-Cd3 at 1:500, rat anti-B220 at 1:500, rat anti-Cd31 at 1:100 (BD). The following day, the sections were rinsed extensively with TBS and incubated with a fluorescent secondary antibody for 5 hours. Secondary antibodies were all used at 1:500 and include donkey anti-rabbit and rat anti-hamster. Sections were washed in TBS then wet-mounted with Vectashield hard set mounting media with DAPI (Vector Labs) and coverslipped.

### Image acquisition

Sections were imaged using 40x, 1.15 numerical aperture and 63x, 1.30 numerical aperture oil objectives on a Leica TCS SPE confocal microscope using Leica Application Suite Advanced Fluorescence software. Stacked images of fluorescent spleen sections were reconstructed using ImageJ (NIH), and Photoshop (Adobe) software was used to change brightness and contrast of the images. In each case all settings were applied equally to each color channel. To quantify double labeling, we used an unbiased approach. For example, to examine whether GFP expression in GFAPcre-GFP mice was present in GFAP+ cells, spleen sections from three animals were immunostained for GFAP with an Alexa Fluor 555-conjugated secondary antibody. When a GFP-expressing cell was identified in the green channel, the channel was switched to red to determine whether it also stained for GFAP. Conversely, to evaluate which percent of GFAP immunostained cells express GFP, GFAP+ cells were identified in the red channel then the channel switched to green to score GFP expression. Each mouse contained three replicates and 100 cells were counted for each of these analyses.

### Dissociation and isolation of spleen glia

Spleens from mice aged 8-12 weeks were removed, minced, resuspended in 4ml of dissociation buffer (HBSS, 10 mM HEPES, 1.4 mg/ml Collagenase A, 0.4 mg/ml DNase I, 5% FBS) and incubated with constant agitation at 37°C for 1 hour. After the hour, Dispase II (2 mg/ml) was added and the samples were incubated for an additional 30 min. Upon completion, 5 mM EDTA was added to the sample and incubated at room temperature for 5 min and subsequently centrifuged at 1000 rpm for 5 min at 4°C. Red blood cells were lysed with 1 ml of ACK buffer (Giboco) for 5 min at room temperature and diluted with 13 ml of cold PBS + 5% FBS before another centrifugation at 1000 rpm for 5 min at 4 °C. The cell pellets were resuspended in 5 ml of PBS + 5% FBS and passed through a 40-micron cell strainer. After washing with 5 ml of PBS + 5% FBS, the remaining cells on the cell-strainer (“size-selected”), were lysed using 500 ul of RLT buffer (Qiagen). RNA from size-selected and whole spleen samples from each animal were purified using the Qiagen RNeasy Micro Plus kit.

### Quantitative RT-PCR

RNA was reverse transcribed using the Invitrogen SuperScript^®^ III First-Strand Synthesis System and used for qPCR using SYBR on the Applied Biosystems™ QuantStudio™ 6 Flex Real-Time PCR System. The standard protocol was used, and all reactions were performed in triplicate with *Gapdh* (F: ACTCTTCCACCTTCGATGCC, R: TGGGATAGGGCCTCTCTTGC) as a housekeeping gene. Primers used were: *S100b* (F: GGTTGCCCTCATTGATGTCTT, R: TTCGTCCAGCGTCTCCAT), *Sox10* (F: CAAGCTCTGGAGGTTGCTG, R: TGTAGTCCGGATGGTCCTTT), and *Ptprc* (F: GGGTTGTTCTGTGCCTTGTT, R: GGATAGATGCTGGCGATGAT). All primers were validated in house using a combination of literature, BLAST and control samples. The ΔΔCt method was used to quantify fold enrichment of size-selected cells over whole spleen samples (Livak & Schmittgen, 2001).

### RNA isolation and sequencing

Cell homogenates from whole spleen and size-selected samples were sequenced from 8 animals (4 male, 4 female) approximately aged 8-12 weeks of age. Quality and quantity of RNA was measured using the Agilent Bioanalyzer PicoChip, and only samples that yielded a RIN number > 9.0 were used. Total RNA (5ng) was then reverse transcribed using the SMART-Seq v4 Ultra Low Input RNA Kit for Sequencing (Clontech), which generates high-quality cDNA from ultra-low amounts of total RNA. Libraries were then constructed from synthesized cDNA using the Nextera XT DNA Library Preparation Kit (Illumina) for RNA-Sequencing. Libraries were sequenced on the Illumina HiSeq 4000 platform to obtain 150bp paired-end reads.

### Mapping and analysis of RNA-sequencing data

Approximately 20-30 million 150bp reads were obtained from all 16 (8 size-selected, 8 whole spleen) samples. These reads were trimmed of adapter sequences, low quality bases, and very short reads using trimGalore! (v0.4.5), a wrapper script which utilizes cutadapt (Martin, 2011) and FastQC (v0.11.7). Remaining reads were aligned to the mouse mm10 genome (GRCm38.p6) using the STAR aligner (Dobin et al., 2013) (v2.7.1), which utilizes an algorithm that minimizes alignment time, mapping errors, and alignment biases. Transcript abundances were then annotated and quantified using the RSEM software package (Li & Dewey, 2011) (v1.3.1). Differential gene expression analysis was then carried out using the DESeq2 (Love et al., 2014) (v1.26) package within the R environment. Transcripts with low abundances (<10 counts for any of the samples) were excluded from analysis. Plots were generated using ggplot2 (v3.2.1).

### Deconvolution and Expression Purification

To determine the proportion of contaminating cells in our size-selected samples, we obtained reference sequencing data from cells likely to be contaminating our samples. Cell type comparison data was obtained from publicly available RNA-sequencing datasets on Gene Expression Omnibus (GEO; https://www.ncbi.nlm.nih.gov/geo/). Cell types and sources can be found in Table 1. Samples were chosen based on data generated from non-transgenic animals who had not undergone any experimental manipulations beyond saline or PBS injections. Gene counts from collected data were used as a reference sample in CiberSortX (Newman et al., 2019), a digital cytometry program which deconvolves cell types based on signature genes generated from reference samples. Signature genes and proportions generated by CiberSortX were used to scale and purify expression in size-selected samples by imputing cell expression from glia samples. The newly generated expression files were used for downstream differential expression analysis.

To define spleen-glia specific genes we used a fold-change cutoff of >3.5 to assure that only genes highly enriched in glia were considered. This cutoff was tested by sensitivity analysis through changing the cutoff ±25% and seeing no changes in the data interpretation.

### Gene expression comparisons

To compare the gene expression of spleen glia to that of other glial cells, gene counts were generated for astrocytes, oligodendrocyte precursor cells (OPCs), oligodendrocytes, enteric glia and spleen glia following the analysis outlined above. Counts were normalized and z-scores were then calculated for each gene and plotted using the ComplexHeatmap Package with default settings in the R environment. Gene clusters were generated using k-means clustering after silhouette computation determined that five clusters were optimal. This partitioning was then used in plotting as a split variable.

### Experimental design and statistical analysis

The aim of this study was two-fold: to establish a complete characterization of spleen glia and identify markers in which hypotheses can be generated for future study. Immunohistochemistry utilized three sections from three animals for a total of nine replicates per stain. Transcriptional profiling was done on eight total animals with four replicates for each sex of both spleen glia and whole spleen samples. Statistical details can be found in the section entitled “Mapping and Analysis of RNA-sequencing”, above. Raw transcriptome datasets generated from size-selected cells and whole spleen were uploaded to GEO (Accession # GSE151856) as both read counts and differential expression tables. To improve visualization of the complete processed data set after deconvolution, a web application was developed; this can be found at *https://buckwalterlab.shinyapps.io/SpleenGlia/*.

## Results

### Spleen glia express S100b and GFAP and ensheath sympathetic axons

We began by defining the relationship between spleen glia, nerves, blood vessels and immune cells in the spleen. To achieve this, we utilized a peripheral glia reporter mouse (C57BL/6J GFAP-cre, ROSA-eGFP) in which pulmonary glia were labelled with GFP (Suarez-Mier & Buckwalter, 2015). To verify that this reporter mouse also expressed GFP in spleen glia, we immunostained spleen sections for two glial markers, GFAP and S100B. We observed GFP-expressing cells with an elongated morphology that co-immunostain for both glial markers in the spleen (Figure 1). The GFP reporter was sensitive and specific: 92 ± 1.5% of GFAP expressing cells expressed GFP, and 99 ± 0.47% of GFP expressing cells co-immunolabeled for GFAP. Additionally, we found that S100B strongly labels spleen glia with high sensitivity, labelling 100% of GFP-expressing cells. The antibody also very weakly labels a subset of macrophages in the spleen, likely due to its known cross-reactivity with S100A6.

**Figure 1:**
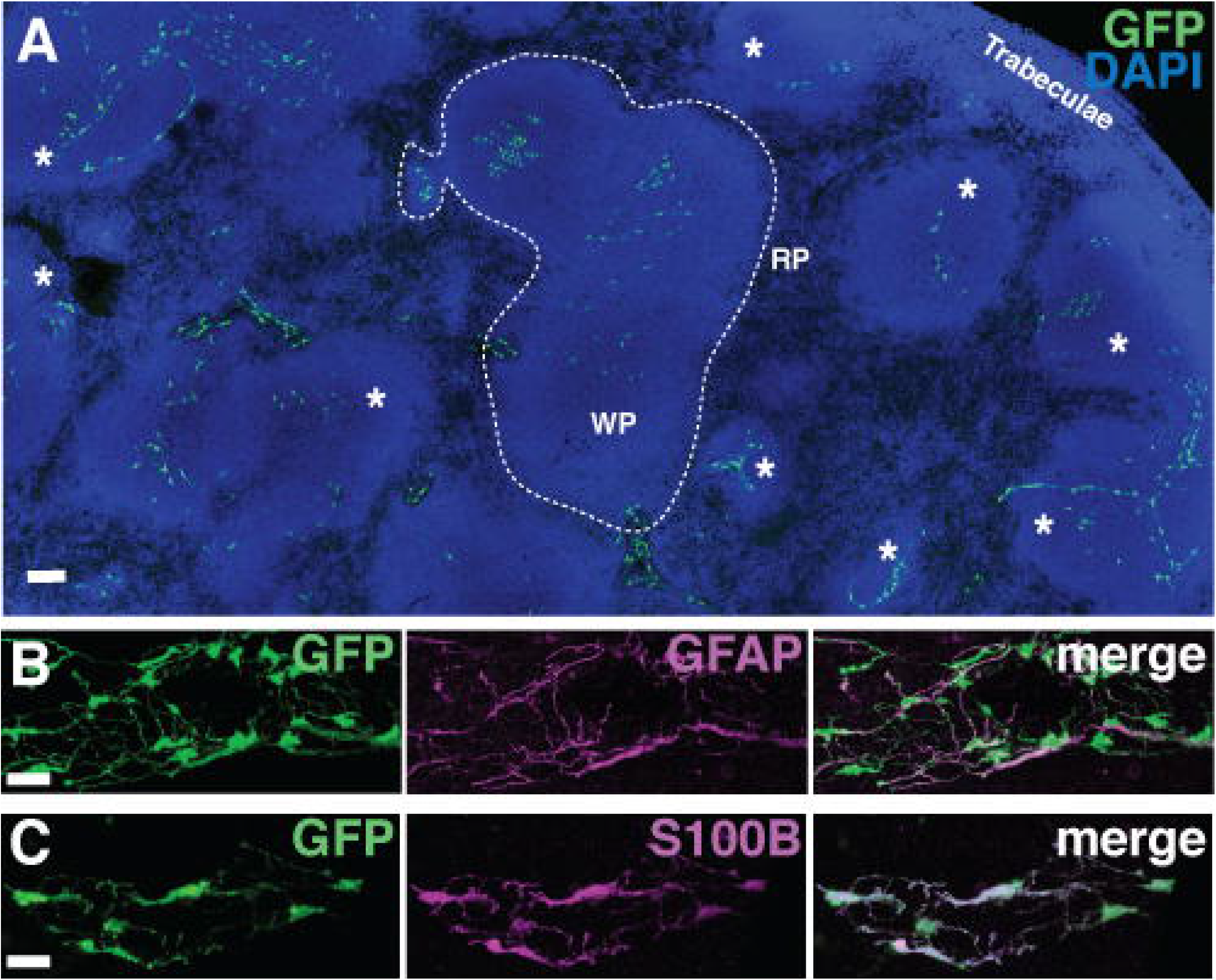
GFAP-cre, Rosa-eGFP reporter mice express GFP in GFAP- and S100B-expressing cells in the spleen. Representative photomicrographs of 40 μm thick spleen sections from GFAP-GFP reporter mice immunostained for (A) GFAP and (B) S100B. Scale bars, 20 μm.

The spleen is innervated by the splenic nerve, which itself is comprised of sympathetic axons that originate from cell bodies in the superior mesenteric ganglion. In general, unmyelinated peripheral nerves can be uniformly associated with glia or have naked terminals (Forbes et al., 1977; Novi, 1968). To assess the association of spleen glia and axons, we immunostained spleen sections for two nerve markers: tyrosine hydroxylase (TH) and protein gene product 9.5 (PGP 9.5). The former is unique to sympathetic and dopaminergic nerves, and the latter is a marker specific to neural and neuroendocrine cells (Figure 2). We observed elongated GFP-expressing spleen glia tightly associated with axons expressing both TH (Figure 2A&B) and PGP 9.5 (Figure 2C&D). Glia tightly intertwined around axons, and we also observed occasional slight axonal protrusions that appear to come to the surface of the glia. These may represent varicosities described in EM experiments where neurotransmitters are postulated to be released through glial fenestrations. Additionally, upon measuring concordance between glia and axons we observed that all glia (100%) were associated with nerve axons while 93 ± 1.2% of axons were associated with glia (data not shown). This result reveals that glia in the spleen are exclusively associated with sympathetic axons, and that nearly all splenic nerve axons are physically associated with glia.

**Figure 2:**
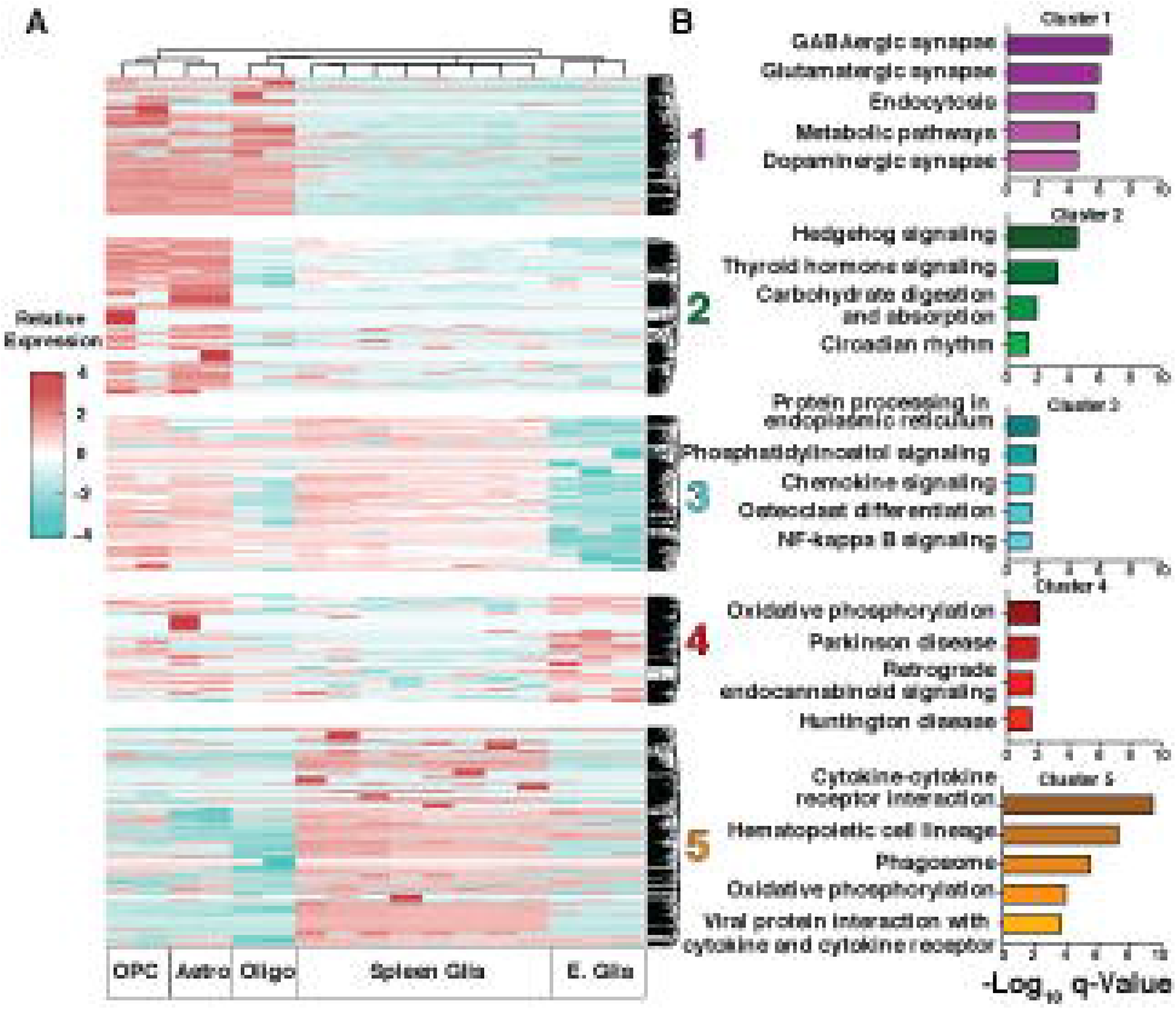
Spleen glia ensheath sympathetic nerves in the spleen. Representative photomicrographs of 40 μm thick sections of spleen. GFP expression is closely associated with nerves expressing tyrosine hydroxylase (TH) (A&B). This close relationship can also be seen using a pan-neuronal marker protein gene product 9.5 (PGP 9.5) (C&D). Both panels show the tight association between nerve and glia. The morphology suggests that axonal protrusions containing neurotransmitter vesicles (TH stain) may protrude out of a fenestrated glial sheath (arrows). Scale bars = 20 μm (A&C), 5 μm (B&D).

### Spleen glia form intricate networks around arterioles that are closely apposed to lymphocytes

The splenic nerve enters the spleen at the hilum, traveling on the outside of the splenic artery. Sympathetic fibers accompany the artery as it branches into fenestrated arterioles that course through the white pulp of the spleen. Fenestrated arterioles have a discontinuous endothelial lining (Aird, 2007), which, where present, can be visualized with the endothelial marker CD31. We observed glia and nerves forming intricate networks around arterioles which were discontinuously lined by CD31-expressing endothelial cells (Figure 3A). To verify that the glia and nerves were located around the splenic arterioles in the white pulp we immunostained spleen sections for T and B lymphocytes (Figure 3B-D). Arterioles were surrounded by a meshwork of glia as they traveled in the peri-arteriolar lymphoid sheath, or PALS region, where CD3_+_ T lymphocytes are housed, and thereafter, as they coursed into B220_+_ B lymphocyte follicles. Thus, there is a close physical relationship between the glia, nerves, and lymphocytes. However, we did not observe similar associations or density when staining for IgM_+_ or Cd1d_+_ marginal zone B-cells (data not shown). Also, the red pulp, which receives its blood supply after arterioles have become capillaries and contains primarily monocytes, macrophages, and red blood cells, exhibited far less staining for either nerves or glia.

**Figure 3.**
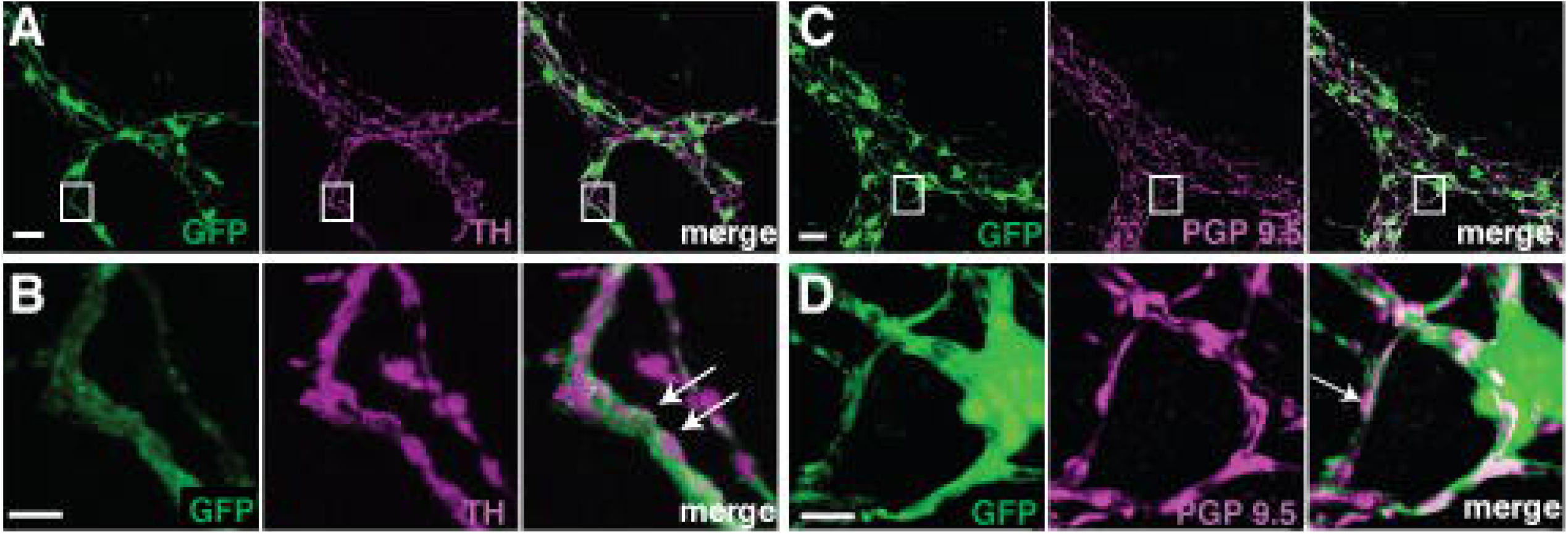
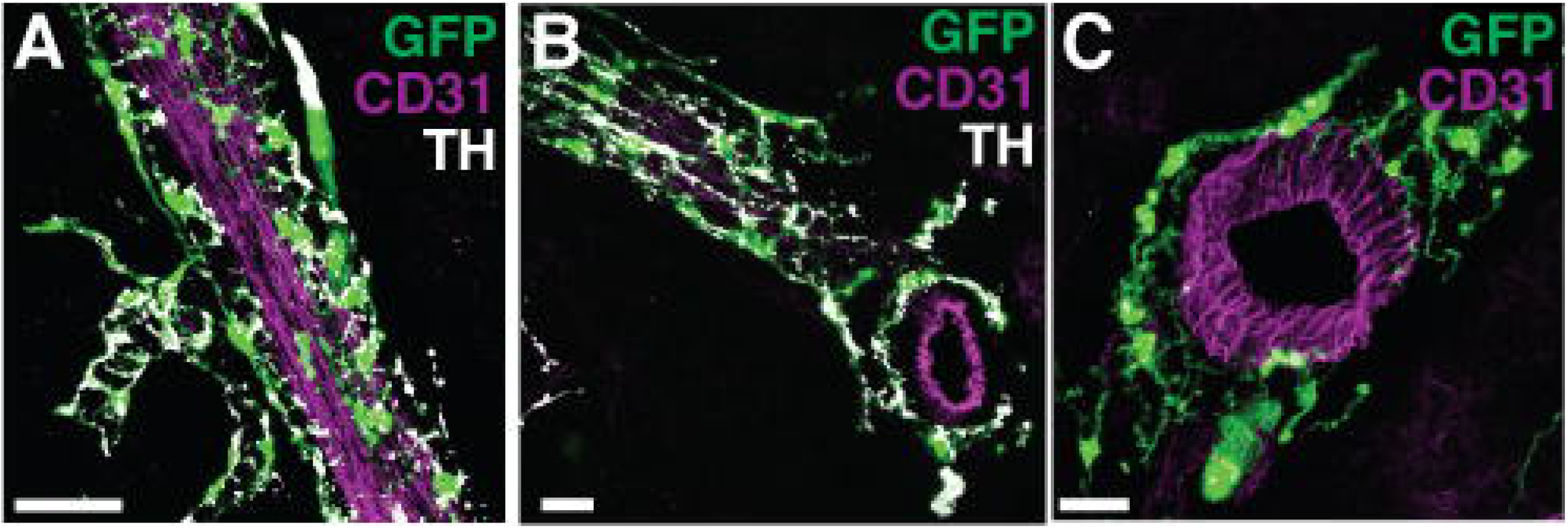
Spleen glia form networks that associate with vasculature as they course through lymphocyte-containing white pulp. Representative photomicrographs of 40 μm sections of spleen depicting thickened processes of spleen glia wrapping around large CD31+ vessels (A). Spleen glia can be seen throughout PALS regions and within B-cell follicles (B&C). The tubular networks of glia branch within B-cell follicles (B), and course through the T-cell-rich PALS regions (D). Scale bars, 20 μm.

### Transcriptional profiling of spleen glia

Our anatomical characterization of spleen glia demonstrates that they are physically juxtaposed between axons and lymphocytes poised to receive molecular signals emanating from the bloodstream (e.g. antigens). This close physical relationship means it is possible that glia participate in neuroimmune signaling and are also likely to provide physical and potentially nutritional support to axons. To determine whether they express genes consistent with these or other functions, we performed RNA sequencing. As described in Materials and Methods, spleen glia were size-selected from spleens of C57BL/6J mice, and total RNA was obtained (Figure 4A) from size-selected spleen glia. Prior to sequencing, RT-qPCR was carried out to confirm expression of known markers of peripheral glia (*S100b* and *Sox10*). Both markers were enriched in the size-selected spleen glia RNA vs. whole spleen RNA, with log2 fold increases of 5.4 ± 0.88, and 5.3 ± 1.2, respectively (Figure 4B). In addition, we observed depletion of protein tyrosine phosphatase, receptor type, C (*Ptprc*) with log2 fold decrease of −1.5 ± −0.65. *Ptprc* is a type I transmembrane protein that is present on all differentiated hematopoietic cells and encodes the immune cell marker CD45. These results indicate that our size selected cells were strongly enriched for glia and de-enriched for immune cells. We selected 8 samples (4 male & 4 female) aged 8-12 weeks that were highly enriched for glial markers *S100b* and *Sox10* (log2 fold enrichment >5) for RNA sequencing analysis.

**Figure 4.**
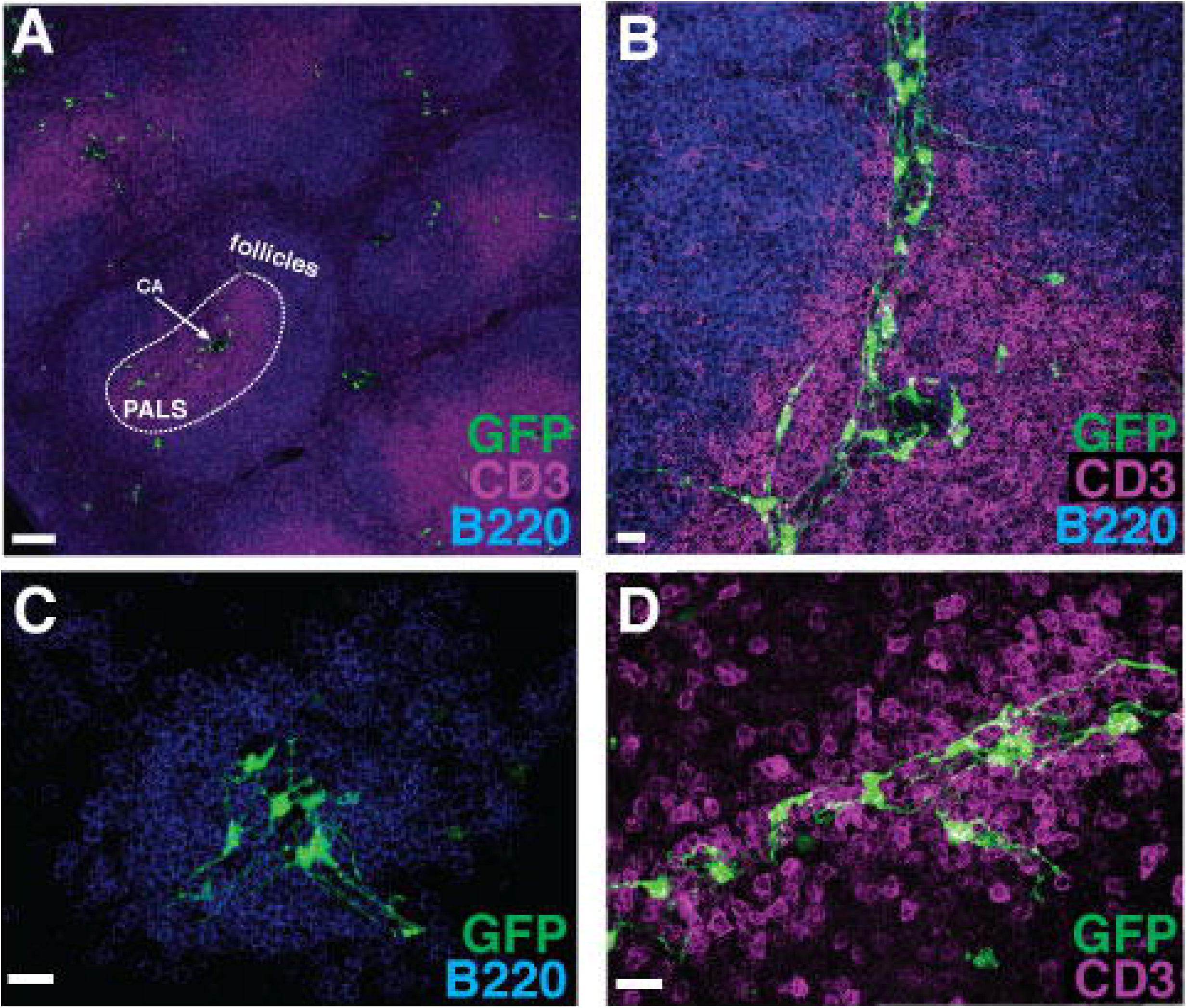
Generating the spleen glia transcriptome. (A) Strategy for isolating and obtaining spleen glia transcriptome and spleen glia enriched gene set. (B) Quantitative real-time PCR of reverse-transcribed genes from RNA selected from size-selected spleen glia prior to RNA sequencing. We observed enrichment of two peripheral glial markers (*SlOOb, SoxlO*) and deenrichment of *Ptprc*, which encodes the cell surface marker CD45 that is expressed on all immune cells. (C) Cell-type deconvolution results after RNA sequencing demonstrate cell-type proportions in bulk samples prior to purification. (D) Enrichment of glial, immune cell, and endothelial cell genes in the spleen glia transcriptome after post-deconvolution purification. Error bars, SEM.

We next utilized cellular deconvolution to further increase specificity for spleen glia genes in our dataset. Deconvolution is a bioinformatic approach that allows researchers to determine the fraction of a cell-type in bulk sequencing data (Kang et al., 2019; Newman et al., 2019). Deconvolution analysis revealed that our sequenced samples contained majority glial cells (mean = 73% ± 4%). All white blood cell types combined accounted for <2% of all genes, while endothelial cells represented the bulk of the contaminating cells, approximately 18% (Figure 4C). Signature gene sets from contaminating cells were then subtracted in proportion to each cell type’s contribution in their corresponding bulk sample, and the newly generated purified spleen glia transcriptomes were used for downstream differential expression analysis.

To define spleen glia-specific genes even more stringently within the purified spleen glia transcriptome, we used an enrichment cutoff of log2 fold >3.5 (FDR <0.01). We refer to this set of stringently selected genes as “spleen glia enriched genes”. Expression cutoff was tested through sensitivity as described in the methods and was determined to be optimal. The log2 fold >3.5 enrichment compared to whole spleen was chosen after we examined fold enrichment of genes common to glia, immune cells and endothelial cells. Differential expression analysis reveals log2 fold enrichment >5 of peripheral glial genes *Sox10* and *S100b* over whole spleen (Figure 4D). Furthermore, there is similar enrichment of other known glial genes including *Atp1b2, Gja1*(*Connexin* 43), and *Plp1*. We then checked for contamination of known markers of immune cells and endothelial cells. Analysis demonstrated that immune cell genes *Ptprc*, T-cell specific (*Cd3*) and B-cell specific (*Cd19*) genes were de-enriched, with log2 fold < −1. Furthermore, we saw only marginal enrichment of endothelial cell specific genes, with log2 fold increases of about 2. We thus chose log2 fold >3.5 for further downstream analysis.

Using the expression cutoffs, we established a “spleen glia enriched genes” list containing 2,202 genes. Table 2 lists the top 25 enriched genes. Several genes on this list are expressed in both PNS and CNS glia, including potassium voltage-gated channels *Kcne4* and *Kcna5*, indicating that spleen glia, like enteric glia and astrocytes, may be involved in homeostatic regulation of extracellular potassium. Moreover, we see robust expression of Neuroligin 3, a cell adhesion molecule involved in maintaining the glial sheath around peripheral nerves (Gilbert et al., 2001), and Tenascin 3, a glycoprotein necessary for gliogenesis (Wiese et al., 2012). Additionally, we examined this gene set for sex differences. However, there were few significant differences (data not shown), so we pooled all samples for the remainder of analysis.

### Spleen glia express various neurotransmitter receptors

Our immunohistochemical characterization demonstrated that spleen glia were always associated with sympathetic axons. Considering this, we next investigated how the nerve and glia might communicate. We searched for genes encoding receptors of molecules known to be released by sympathetic axons including norepinephrine (NE), and to a lesser extent epinephrine (EPI), purines, and neuropeptide Y. We examined all receptors for these three neurotransmitters that were present in the whole spleen and the spleen glia transcriptome (Figure 5A). We indeed observed significant expression and enrichment of various adrenergic receptors in spleen glia compared to whole spleen. Of the adrenergic receptor subtypes, the β2-adrenergic receptor was most highly expressed by spleen glia. The β2-adrenergic receptor is also expressed by astrocytes, enteric glia, and Schwann cells. In addition, our data showed that spleen glia highly express α-adrenergic receptors, three of which are highly enriched compared to whole spleen (*Adra1b, Adra1d*, and *Adra2c*). This is notable because the α1-adrenergic receptor has a higher affinity for NE than EPI (Rang et al., 2016) and the splenic nerve predominantly releases NE (Kirpekar & Misu, 1967).

**Figure 5.**
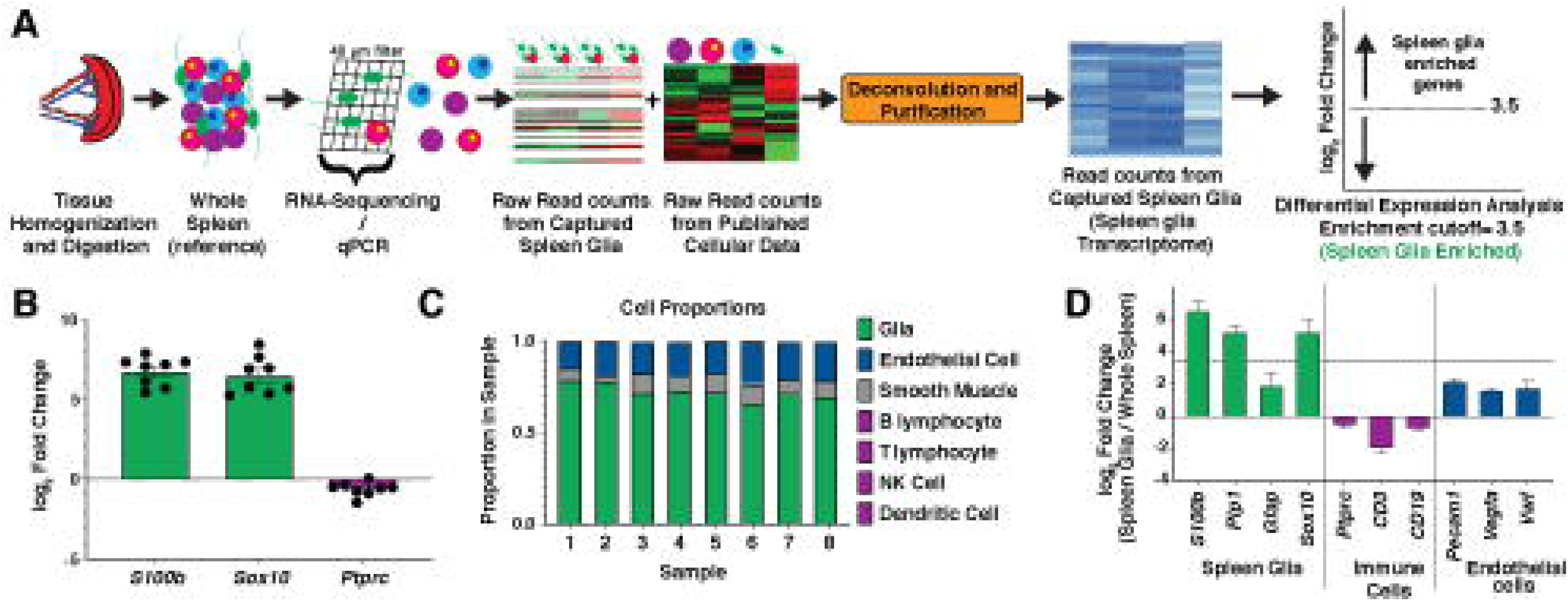
Heatmaps of neurotransmitter receptors genes expressed by spleen glia. Boxes to the right of each gene are colored according to expression level in spleen glia (left) and whole spleen (right). Bolded gene names indicate genes that are highly enriched in spleen glia compared to whole spleen (log_2_FC > 3.5, TPM > 1, FDR < 0.01). Neurotransmitter receptor genes known to encode receptors for sympathetic nerve messengers. Spleen glia express neurotransmitter receptors for norepinephrine (adrenergic receptors), neuropeptide Y, and ATP. TPM, transcripts per million; FDR, false discovery rate.

Spleen glia were also more enriched for Neuropeptide Y receptor 1 expression than whole spleen RNA. Neuropeptide Y is primarily released by sympathetic nerves in the PNS (Tan et al., 2018). Expression of its receptor in Schwann cells has already been validated (Park et. al 2015). Its exact role, however, has not been fully elucidated.

Spleen glia also enrich for several purinergic receptors (Figure 5A). ATP has long been recognized as an intracellular energy source, but it is also a potent extracellular neurotransmitter co-released by sympathetic nerves (Westfall et al., 2002). Purinergic signaling in glia has been linked to glial proliferation, motility, and survival as well as cytokine signaling and injury responses in Schwann cells and astrocytes. *P2ry2* was the most highly enriched purinergic receptor subtype. This subtype has known functions in astrocytes inducing activation and release of inflammatory molecules (Erb & Weisman, 2012).

### Pathway analysis reveals expected and potentially novel roles for spleen glia

Spleen glia likely serve similar functions as other peripheral glia, such as ensheathing and supporting nerves, guiding axons during axon outgrowth and response to injury and disease. However, they may serve other functions as well. To assess their likely functions in an unbiased fashion, we carried out pathway analysis on the spleen glia enriched gene list (TPM>1; log2 FC >3.5; FDR < 0.01) (Figure 6).

**Figure 6.**
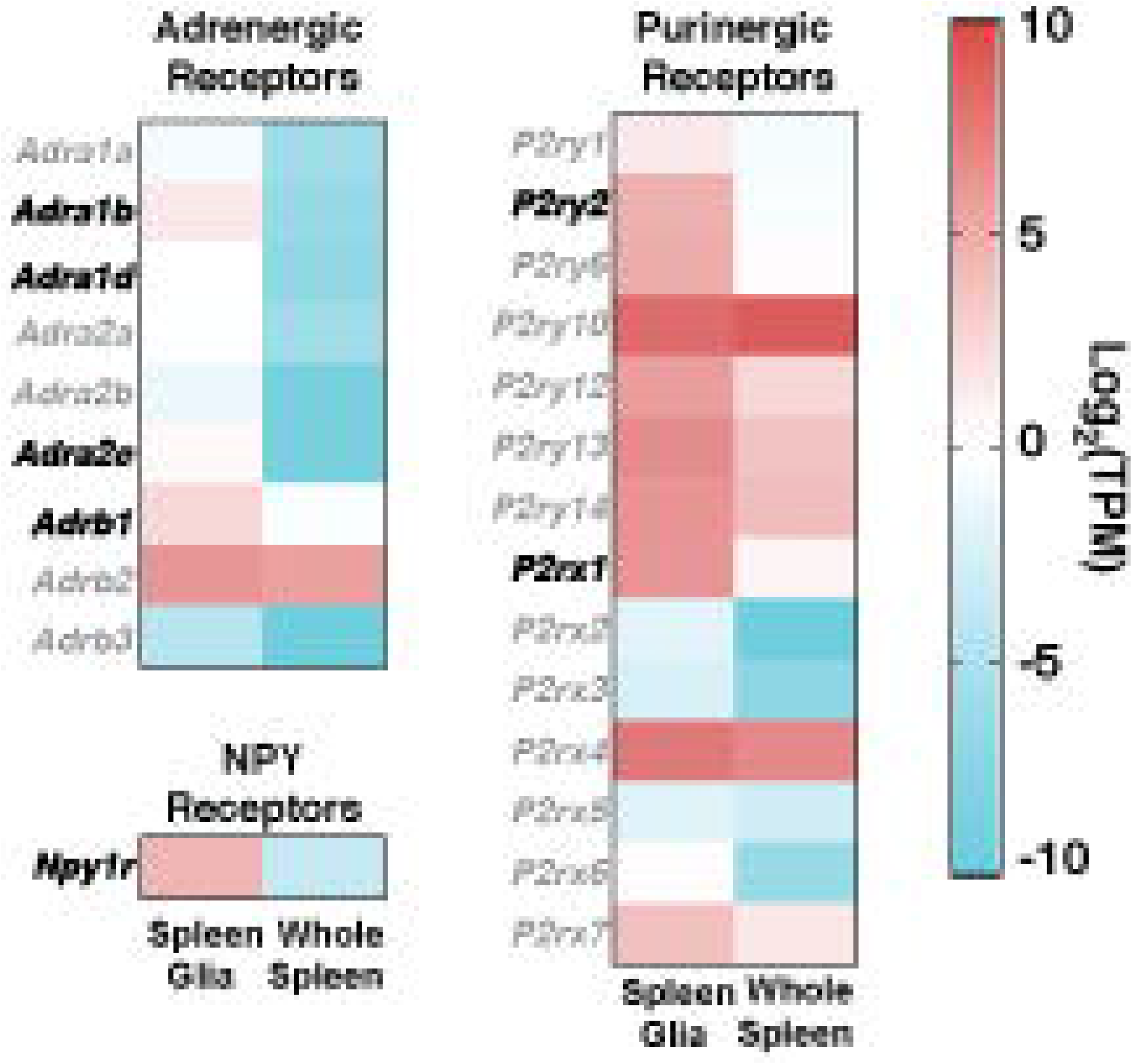
Pathway analysis of significantly enriched spleen glia genes reveals roles in focal adhesion, axon guidance, and immune responses. (A) Bar graph representing the top 10 KEGG Pathways in spleen glia. Numbers to the right of bars indicate the number of genes enriched in glia / total gene number in the pathway. (B-F) Heatmaps of genes in these KEGG pathways that are enriched in spleen glia (left) with expression in whole spleen (right), colored by expression level in transcripts per million (TPM, see scale). (B) The Focal Adhesion KEGG pathway contains thrombospondins, laminins, and Erb-B receptors. (C) Selected genes from the Axon Guidance KEGG pathway, many with known roles in axonal support by other peripheral glia cell types. (D-F) Selected genes from the immune-related KEGG pathways–AGE-RAGE signaling, Cytokine-cytokine receptor interaction, and Complement cascade. All genes used in KEGG analysis are from the spleen glia enriched gene set (log2 FC> 3.5, TPM>1, FDR<0.01 compared to whole spleen).

Interestingly, one of the most highly enriched pathways was the focal adhesion pathway (Figure 6A). Spleen glia highly expressed several known focal adhesion genes such as thrombospondins and laminins (Figure 6B). Both molecules have known functions in other glial cell types and activate downstream Pi3k-Akt signaling, which is also enriched in spleen glia. For example, thrombospondin-2 expression in astrocytes is necessary to regulate synaptic adhesion (Christopherson et al., 2005). In addition, spleen glia express ErbB receptors, which suggests they are active participants in NRG1-ErbB2/3 signaling, a fundamental cell adhesion signaling pathway between Schwann cells and axons (Newbern & Birchmeier, 2010). Another known function of Schwann cells is facilitation of axon guidance, and pathway analysis also determined that this process is significantly enriched in spleen glia. Genes for ephrins, semaphorins, and plexins were extensively represented (Figure 6C). Members of these gene families are described in so-called “repair” Schwann cells that promote axonal regrowth after injury (Koncina et al., 2007).

Glia may have organ specific functions, and spleen glia reside in an organ that is heavily involved in immunity. Accordingly, we hypothesized that spleen glia may be active partners in mediating immune responses. Indeed, AGE-RAGE signaling, cytokine-cytokine receptor interaction and complement cascade signaling were among the top 10 enriched pathways (Figure 6). Each of these pathways are driven by expression of immune relevant genes including cytokines and chemokines, interleukins and their receptors, and complement proteins. Interestingly, although some cytokines and chemokines are enriched in spleen glia, many of the most enriched genes are receptors. For example, the gene for Gp130 (*Il6st*) is enriched, and critical for astrocyte activation and ability to control infection in the brain (Drögemüller et al., 2008). Gp130 is a receptor that associates with a number of cytokine receptors to transmit signals to JAK/STAT3 or 5. Spleen glia express many of these gp130 co-receptors, including *Osmr, Cntfr, Il6ra, and Lifr*, implying that spleen glia likely respond to oncostatin and/or IL31, Cntfr, IL6, and LIF, respectively. Several transcripts for cytokines and chemokines including *Il34, Il7*, and *Cxcl13* are also enriched in spleen glia, as are the TBGβ family members *Tgfb2, Gdf6* and *Gdf10*.

### Spleen glia exhibit similarities and differences compared to other glia

To elucidate differences between glial cell types, we compared the spleen glia transcriptome to publicly available transcriptomes of astrocytes, oligodendrocyte precursor cells (OPCs), oligodendrocytes, and enteric glia outlined in Table 1. *Z* scores were calculated for each gene, plotted on a heatmap, and cell types were arranged after unbiased hierarchal clustering analysis (Figure 7A). Spleen glia shared many genes with all these glial cell types and were most similar to enteric glia. Clustering analysis identified five unique gene groupings across all the cell types. We performed pathway analysis on the genes in each cluster (Figure 7B).

**Figure 7.**
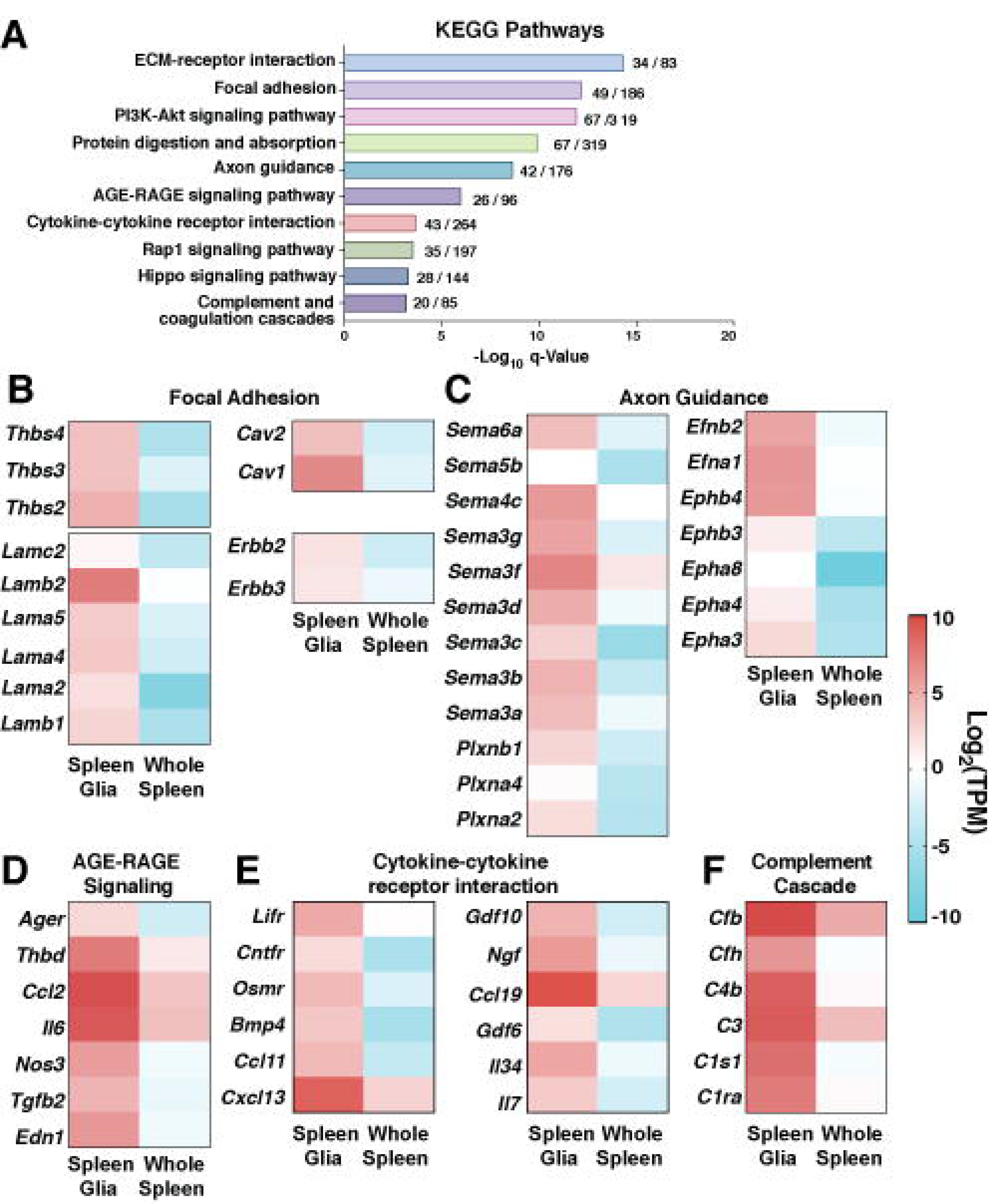
Transcriptome comparison between spleen glia and other glial types. (A) *Z*-score Heatmap of transcriptomic data from five glial types after hierarchical clustering analysis identified five main clusters. Each row is one gene, and color represents relative expression of that gene in the cell type of its column compared to its expression levels in all other cell types. (B) Bar graphs representing top KEGG Pathways from each cluster. OPC, oligodendrocyte precursor cell; Astro, Astrocyte; Oligo, Oligodendrocyte; E. Glia, Enteric Glia.

Cluster 1 was highly expressed in CNS glia compared to PNS glia. The pathways enriched in this cluster represent GABAergic, glutamatergic and dopaminergic synapse signaling, all present in the CNS but perhaps not dominant in the PNS. Cluster 2 represented genes enriched in OPCs and astrocytes but not oligodendrocytes. Hedgehog signaling was the most abundant pathway of this cluster and has known roles in both cell types. It is involved in the formation of mature oligodendrocytes from OPCs, and the regulation of inflammation by astrocytes (Allahyari et al., 2019; Laouarem & Traiffort, 2018).

Cluster 3 represented genes that are downregulated in enteric glia compared to spleen glia and Cluster 4 was upregulated in enteric glia compared to spleen glia. Pathways were least significant for these two clusters, possibly due to more shared genes and functions between all cell types. However, Cluster 3, composed of the genes with the lowest expression in enteric glia, contained immune genes found in spleen glia as well as other glial cell types, including those in the “chemokine signaling” and “NF-κb signaling” pathways. Both pathways suggest a role for spleen glia in immune regulation. There is also extensive evidence that OPCs and astrocytes use chemokines to regulate myelination and inflammation in the CNS. Genes in Cluster 4 were generally lower in spleen glia and predominately enriched in enteric glia. The top pathway here was “oxidative phosphorylation,” a process which ultimately leads to the production of ATP.

Cluster 5 contained genes that were uniquely enriched in spleen glia over other glial cell types. The top enriched pathway within this cluster was “cytokine-cytokine receptor interactions,” which included genes encoding cytokine and chemokine receptors and their respective ligands. This finding suggests that spleen glia may be more involved in initiating and responding to inflammation than other glial types. Additionally, Cluster 5 contains genes in the phagosome pathway, which is notable because phagocytosis is a central process in inflammation and defense against virus and bacteria. Expression of genes for the toll-like receptor gene *Tlr2* and antigen processing gene *Tap1* specifically seem to suggest the utilization of toll-like receptor promotion of ER-mediated phagocytosis (Desjardins, 2003).

This is consistent with pathway analysis on the enriched spleen glial genes alone and implies that spleen glia may be directly involved in both innate and adaptive immune responses. Although further research needs to be done to determine exact mechanisms, we have demonstrated that spleen glia are anatomically and genetically poised to participate in immune responses in the spleen. The processed spleen glia transcriptome, after deconvolution, is available as a resource for future study (https://buckwalterlab.shinyapps.io/SpleenGlia/).

## Discussion

Here we provide an in-depth anatomical and transcriptional characterization of spleen glia, while identifying a mouse reporter as a tool to study them. Spleen glia are elongated cells visualized using immunostaining for GFAP and S100B. They maintain a tight association with nerves in white pulp regions and come into close contact with lymphocytes. We identified genes involved in the ensheathment of axons and communication with both nerves and immune cells. Spleen glia express many genes in common with other glial types; the predominant difference is that spleen glia express immune genes more highly. This transcriptome is a first step towards better understanding spleen glia and implies that spleen glia are unlikely to function solely as support cells for axons. Rather, they are most likely active players in neuroimmune communication and function.

Spleen glia have a uniform morphology forming dense networks that ensheath sympathetic axons. Another study describing immunohistochemical localization of spleen glia (Ma et al., 2018) used a GFAP antibody that exhibited more widespread staining than we observe. In addition to the cells we observe, they observe extensive staining throughout the red pulp and marginal zone that is not present in our reporter mice, or our GFAP or S100B stains, and is not adjacent to axons. Because we don’t see this staining with any of our methods, we believe that their immunostaining reflects some background contamination.

Interestingly, the axonal morphology we observe at higher power appears to show protruding varicosities from axons with less glial coverage at these sites. This is similar in appearance to axon varicosities described in literature, where glia form fenestrations at sites of neurotransmitter release (Burnstock, 2008; Douglas & Ritchie, 1962). In this case, the target cells for sympathetic neurotransmitters are likely immune cells, glia, and vascular smooth muscle cells.

Glial ensheathment of axons is likely maintained through several molecular signaling pathways. For instance, spleen glia express ErbB receptors, important for the Neuregulin 1 (Nrg1)-ErbB2/3 receptor cascade that facilitates Schwann cell-axon interactions. Neurons produce Nrg1 which interacts with ErbB2/ErbB3 receptors on glia. This is critical for Remak cells, a type of non-myelinating Schwann cell ensheathing small axons such as C-fibers (Harty & Monk, 2017). Loss of Nrg1 in sensory axons results in more axons per Remak bundle and reduced Remak cell ensheathment. This pathway likely functions similarly in spleen glia, determining which and how many axons become ensheathed. Spleen glia also express Neuroligins, a family of cell adhesion proteins expressed in other glial types. Neuroligin 3 is a vertebrate gliotactin necessary for the development of the glial sheath in the PNS of *Drosophila melanogaster* (Gilbert et al., 2001). Furthermore, loss of astrocytic Neuroligin 2 leads to a decrease in both astrocyte size and cortical excitatory synapses (Stogsdill et al., 2017). Neuroligins likely also function similarly in spleen glia to tether and ensheath axons.

Spleen glia and sympathetic axons course around arteries and arterioles in the spleen, typical for sympathetic innervation throughout the body. In the spleen, lymphocyte-rich white pulp regions surround arteries. The anatomical location of spleen glia suggests involvement in immune responses. Pathway analysis supports this as well. We carefully considered whether our spleen glia transcriptome is contaminated by genes from immune cells in the spleen. Although we cannot eliminate the possibility of any contamination by immune cells, our initial results on size-selected cells prior to deconvolution and purification indicates low contamination of immune cells (<2%). There is also de-enrichment of cognate immune cell genes for lymphocytes, white blood cells, MHC class II, and immunoglobulin genes. We also considered whether there was bleed-through of cytokine and chemokine gene expression from the whole spleen. Of the top five expressed chemokines and cytokines in whole spleen, four are de-enriched in spleen glia (*Ccl5, Ccl4, Tgfb1*, and *Ccl6*). The remaining one, *Cxcl1*, didn’t meet our cutoff of enrichment (log2fold = 2.6) in spleen glia but is also expressed by both astrocytes and Schwann cells (Ni et al., 2019; Ntogwa et al., 2020).

More work will be needed to elucidate the true role of spleen glia in immune responses. However, immune gene expression is not unique to spleen glia; for example, astrocytes and Schwann cells also express complement proteins (de Jonge et al., 2004; Lian et al., 2016). The complement cascade functions to initiate and propagate inflammation. Spleen glia may participate in the alternative complement pathway, as they differentially express complement factors Factor B and H (Hoffman & O’Shea, 1999). Moreover, spleen glia express pro-inflammatory cytokines that are upregulated during complement activation, such as *Nos* and *Il6*, both expressed in astrocytes (Hamby et al., 2006; Van Wagoner et al., 1999). Since we generated our spleen glia transcriptome in the absence of immune stimulation, the expression of these cytokines suggests they serve homeostatic immune functions. Our data also demonstrate high expression of *Il15* by spleen glia (TPM=11.8), which promotes long-term proliferation of activated T-cells (Sprent et al., 2000).

A limitation of this study is our isolation protocol. Due to the size, shape and density of spleen glia, common RNA isolation protocols produce RNA at lower quality and quantity than what is needed for next-generation sequencing. In developing our isolation protocol, we found that the cells are rare, that RNAses are high in the spleen (limiting the usefulness of RiboTag ribosomal pulldown methods), and that FACS sorting pulverizes the cells. However, we took advantage of the large size of spleen glia to develop a size-selection method that yielded sufficient quantity and purity of glial RNA. We utilized a stringent bioinformatic approach including cellular deconvolution and expression cutoffs to maximize the specificity of our gene lists. A notable casualty of this approach was *Gfap*. Although the *Gfap* promotor was effective at labeling spleen glia for anatomical characterization, we discovered that its RNA expression level was too low to make it a reliable genetic marker. This is consistent with enteric glia literature that describes *Gfap* expression to be lower than other glial markers such as *S100b* (Rao, Nelms, Dong, Salinas-Rios, et al., 2015). This may indicate that, as in the brain, GFAP is a highly stable protein that is upregulated during glial proliferation and development and remains for long periods of time without robust gene expression (Rolland et al., 1990). There are likely other casualties for similar reasons, however our final transcriptome errs on the side of including only genes that are highly enriched in spleen glia.

Spleen glia expressed many genes in common with other glial cell types, including receptors for neurotransmitters that facilitate communication with neurons and nerves, including purinergic and adrenergic receptors (Lecca et al., 2012). In sympathetic nerves, ATP is released as a co-transmitter alongside NE. Signaling through purinergic receptors induces various responses in glial cells depending on receptor subtype and context. For example, signaling through P2X ion channels induces a pro-inflammatory response in enteric glia and astrocytes (Bhave et al., 2017; Gandelman et al., 2010), while signaling through P2Y G-protein coupled receptors induces proliferation of astrocytes (Quintas et al., 2011) and calcium responses in enteric glia (Brown et al., 2016). Spleen glia express both subtypes and the necessary machinery for calcium signaling, including the gap junction gene connexin 43, suggesting that they may similarly have calcium waves induced by purinergic signaling.

Adrenergic signaling in glia affects inflammation, behavior, and disease, most studied in astrocytes. For example, β2 adrenergic signaling in astrocytes initiates TNF induced inflammation (Laureys et al., 2014), while α2 receptors stimulate GABA release to suppress neuronal activity (Gaidin et al., 2020). Spleen glia highly express both subtypes, with α1 receptors being the most differentially expressed. Adrenergic signaling in the spleen is important in immune cell activation and proliferation (Bai et al., 2011; Madden et al., 1994). As in astrocytes, adrenergic signaling in spleen glia likely plays a role in altering transmission to target cells such as immune cells. Since many immune cells also express adrenergic receptors, cell-specific knockout experiments will help us understand their role in spleen glia.

This work is an essential first step towards understanding neuroimmune communication in the spleen. We demonstrated that spleen glia form extensive networks throughout the white pulp of the spleen, are transcriptionally unique, and are likely involved in neuroimmune communication. One question that interests us is whether spleen glia are like non-myelinating Schwann cells in other secondary lymphoid organs such as lymph nodes and Peyer’s Patches, and another is how they resemble other visceral glia. Our data also provides a starting point to explore what changes occur in spleen glia during active immune responses, and whether they regulate neuroimmune communication. The reporter mouse used in this study and the web application we created for exploring the transcriptome data provide critical tools to aid future research into spleen and other visceral glia.

## Acknowledgments

This work was funded by the National Institutes of Health, R21 NS098716 to MSB and Illumina Core lab grant, NSF GRFP DGE-114747, Stanford VPGE DARE fellowship to TAL. We also would like to acknowledge Kendra Joan Lechtenberg for her help with manuscript revisions.

## Conflict of interest statement

The authors declare no competing financial interests.

---

**Table 1.** Cell transcriptomes used for deconvolution, and comparison to spleen glia.

**Table 2.** Top 25 most differentially expressed genes in spleen glia. Log2 fold change is relative to whole spleen. FC, fold change; FDR, false discovery rate; TPM, transcripts per million reads.

## Notes

### Competing Interest Statement

The authors have declared no competing interest.

